# Estimating a novel stochastic model for within-field disease dynamics of banana bunchy top virus via approximate Bayesian computation

**DOI:** 10.1101/762765

**Authors:** Abhishek Varghese, Christopher Drovandi, Kerrie Mengersen, Antonietta Mira

**Affiliations:** School of Mathematical Sciences, Queensland University of Technology, Brisbane, Australia; ARC Centre for Excellence in Mathematical and Statistical Frontiers (ACEMS); Institute of Computational Science, Università della Svizzera italiana, Lugano, Switzerland; Department of Science and High Technology, Università degli Studi dell’Insubria, Como, Italy

## Abstract

The Banana Bunchy Top Virus (BBTV) is one of the most economically important vector-borne banana diseases throughout the Asia-Pacific Basin and presents a significant challenge to the agricultural sector. Current models of BBTV are largely deterministic, limited by an incomplete understanding of interactions in complex natural systems, and the appropriate identification of parameters. A stochastic network-based Susceptible-Infected model has been created which simulates the spread of BBTV across the subsections of a banana plantation, parameterising nodal recovery, neighbouring and distant infectivity across summer and winter. Findings from posterior results achieved through Markov Chain Monte Carlo approach to approximate Bayesian computation suggest seasonality in all parameters, which are influenced by correlated changes in inspection accuracy, temperatures and aphid activity. This paper demonstrates how the model may be used for monitoring and forecasting of various disease management strategies to support policy-level decision making.

**Author summary:** The Banana Bunchy Top Virus (BBTV) poses one of the greatest threats to the food security of developing nations and the banana industry throughout the Asia-Pacific Basin. Decision-makers face significant challenges in mitigating BBTV spread in banana plantations due to the vector-borne spread of this disease, which is significantly influenced by a vast array of external environmental factors that are unique to each plantation.

We propose a flexible network-based model that describes the spread of BBTV in a real banana plantation through a random process while accounting for individual plantation characteristics and utilise a principled methodology for estimating model parameters. Our findings quantify the effect of seasonal changes and plantation configuration on BBTV spread and predict for high-risk areas in this plantation. We believe that our model might be used by decision-makers to evaluate the effectiveness of current disease management strategies and explore opportunities for improvements.

## Introduction

The Banana Bunchy Top Virus (BBTV) is one of the most economically important vector-borne banana diseases throughout the Asia-Pacific Basin. The disease was first introduced to Australia in 1913 via infected suckers from Fiji, and spread locally through the banana aphid, *Pentalonia nigronervosa* [1]. With limited knowledge on epidemiological characteristics of the disease or disease management approaches, incidence rates across Australian banana plantations rose rapidly, eradicating over 90% of national crop production in the 1930’s [2]. Cook et al. [3] estimate that the economic benefits of BBTV exclusion from commercial plantations range from $15.9 to $20.7 million each year, approximately 5% of annual crop production value. Aggressive disease management strategies implemented by the Australian Government from the 1930s-90s have largely restricted the disease to the south-east Queensland and northern New South Wales regions of Australia [4]. Eradication, however, has not been achieved, requiring continuous monitoring by the National BBTV Program.

While monitoring the infection counts across the region does provide an indication of disease management success, a vast array of external environmental factors influence BBTV growth, making this an unreliable metric. Furthermore, there are currently limited opportunities to explore various management strategies for BBTV within banana plantations, which could reduce infection rates and identify cost-saving measures for the monitoring program. In such scenarios, mathematical models offer the opportunity to simulate various disease management strategies with a low-cost and quick turnaround [5].

Unfortunately, there have been few contributions to modelling the disease dynamics of BBTV in plantations in the last few decades – despite the significant advancements in computational resources and our understanding of vector-borne diseases. Allen [6] generated a stochastic spatiotemporal polycyclic model for BBTV to describe disease progress within a banana plantation, specifically focusing on identifying the mean inoculation distance of BBTV. However, the model was designed for a hypothetical homogenous circular plantation, which does not account for the various plantation configurations and unique plantation characteristics present around the world; a key factor which greatly affects the effectiveness of disease management strategies. Another model developed by Smith et al. [1] aimed to describe the influence of external inoculum on BBTV spread within a banana plantation in the Philippines. However, the model developed by Smith et al. [1] is deterministic, which results in poor representations of complex natural processes that are inherently probabilistic. Furthermore, these articles do not provide a principled parameter estimation method with appropriate uncertainty quantification based on available field data.

In our paper, we propose a new stochastic model that describes BBTV spread across a banana plantation, parameterising for neighbouring and long-distance infectivity rates, and the subsection recovery rates. Further, we develop a principled Bayesian parameter estimation method for calibrating this model to real field data. Given the intractability of the likelihood function for this model, we employ approximate Bayesian computation (ABC) [7] for estimating model parameters and their uncertainty. Our methodology is inspired by Dutta et al. [8], who demonstrate that ABC may be effectively used to estimate the spreading parameters of a disease by applying a simple Susceptible-Infected (SI) model over a known network structure. This paper adapts and extends this approach to further understand the spreading characteristics of BBTV, evaluate various disease management strategies at the plantation level, and predict the spread of future outbreaks. Even though our work is motivated by the vector-borne transmission of BBTV, we believe that our modelling framework is easily adaptable to describe the within-field disease dynamics of other vector-borne diseases.

## Methods

This study focuses on a banana plantation in Newrybar, near the North-Eastern border of New South Wales in Australia (28°42’14.8”S, 153°32’20.4”E), with an area of approximately 12 hectares. A routine site inspection in 2013 identified a banana plant infected with BBTV in the north-western region of the farm. A following inspection in 2014 identified 26 infections clustered across the South-Eastern region of the farm. Since 2014, the site has undergone monthly inspections, collecting location data and plant characteristics, while implementing a rogue-and-remove disease management strategy. The location of every infected plant has been recorded using the Global Positioning System (GPS) functionality on a smartphone. The dataset consists of 38 months of infection data with the coordinates of each infected plant identified and rogued during a site visit to the farm.

Fig 1 provides a birds-eye view of the plantation. The plantation is separated by dirt paths approximately 3-5 m wide, creating smaller rectangular subsections of banana plants across the plantation.

**Fig 1.**
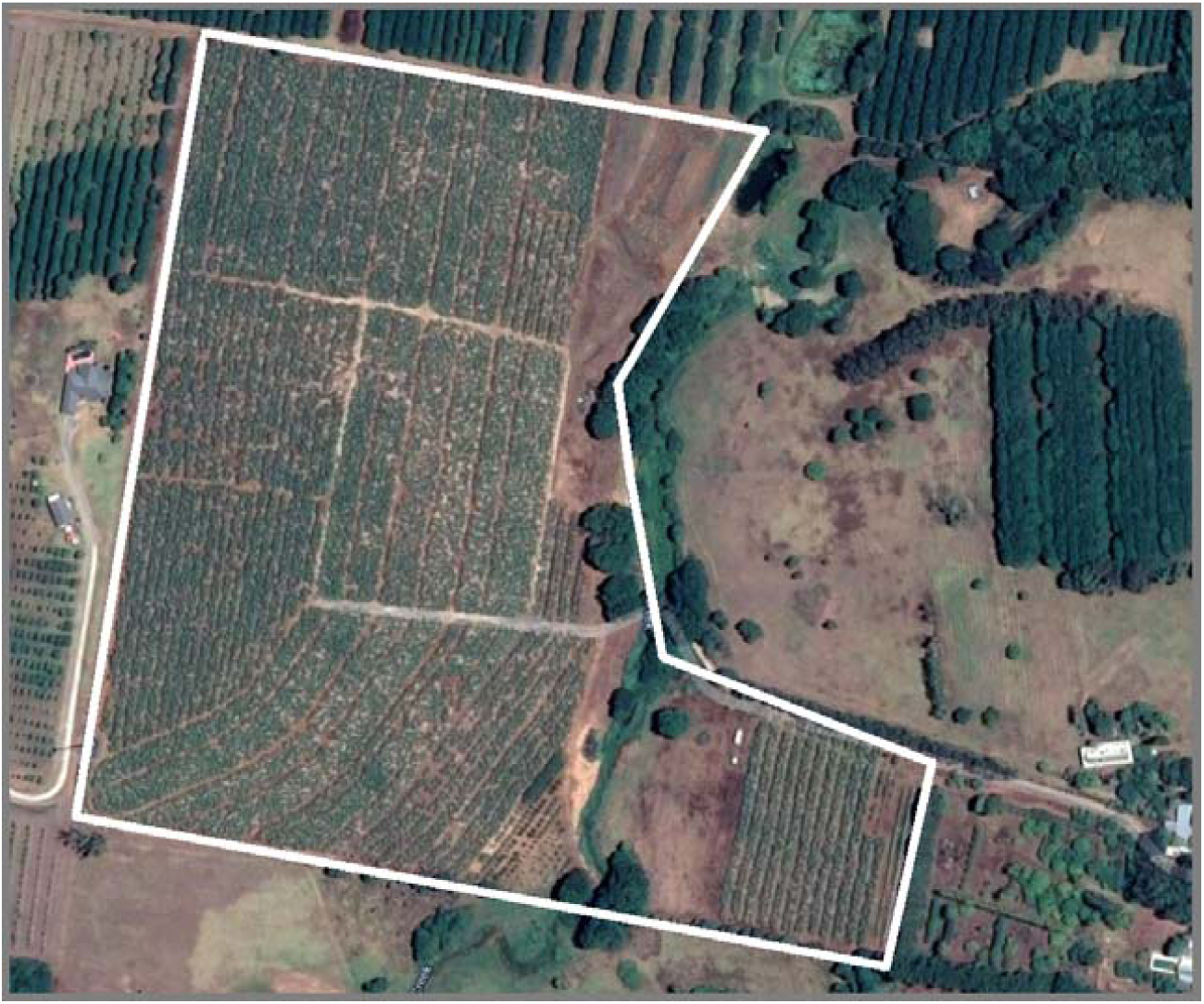
Satellite Image of Newrybar Banana Plantation. Solid white lines indicate the approximate border of the property.

### BBTV Forward-Simulation Model

We propose to model the spread of BBTV in a banana plantation by modifying the ‘simple contagion’ model developed by Dutta et al. [8]. The ‘simple contagion’ model simulates a standard SI process on a fixed network structure. At each time step, each infected node chooses one of its neighbours with equal probability regardless of their status (susceptible or infected), and if the chosen node is susceptible, it is infected with probability θ. Our network-based forward-simulating modelling approach enables easy adaptation to describe the disease dynamics over a range plantation structures, and various vector-borne diseases.

Dutta et al. [8] denote this model by *M*_*s*_ and parameterise it in terms of the spreading rate θ and the seed node *n*_*sn*_. For given values of these two parameters, *n*_*sn*_ = *n*_*sn*_*\** and θ = θ*, they forward simulate the evolving epidemic over time using the model *M*_*s*_.

We adapt the model *M*_*s*_ in several ways for our application. Dutta et al. [8] determine a node to be an individual person, with edges representing each person’s contacts. Modelling the individual status of every plant as a node would be impractical and computationally expensive, due to the varying contact points between plants, and the complex plant-pathogen-vector relationships as described in the previous section. Instead, the data may be aggregated to monitor the infection likelihood over larger areas of the farm. The wide dirt paths throughout a farm act as a soft barrier to BBTV spread, since most aphids are apterous (wingless) and are likely to move from leaf-to-leaf. Therefore, the subsections in a plantation may be described as nodes, where any subsection containing at least one infected plant may be considered ‘infected’.

Secondly, *M*_*s*_ estimates for a single seed node which propagates infection spread in the network. This may be an unrealistic assumption for modelling BBTV spread in plantations, since it is possible for a plantation to have multiple latent infections upon first exposure to the virus. Therefore, *M*_*s*_ must be adapted to accept multiple seed nodes. Additionally, since the scope of this paper is limited to the analysis of current trends and predictions to evaluate disease management strategies, the new BBTV model does not estimate for an initial seed node. Rather, the BBTV model considers the infected nodes observed in the first month of field data surveying to be the seed nodes.

#### Parameters

The unique epidemiological characteristics of BBTV must be parameterised to fully capture the complex dynamics of BBTV transmission in plantations, and to extend on the current literature on BBTV.

There are three parameters that are estimated in this model:

1. *Probability of recovery, θ*_*0*_: As infected plants are rogued and removed at monthly inspections of infected plantation visits, nodes may subsequently recover from an infected state to a susceptible state. Currently, there is no existing literature on BBTV recovery rates in field scenarios using current disease management strategies. Accurate estimates of the probability of node recovery in plantations could inform decision makers on the effectiveness of current rogueing methods, inspection frequency and inspection accuracy.
2. *Neighbouring probability of infection, θ*_*1*_: The model *M*_*s*_ created by Dutta et. al [8] proposes an infection probability for each of the neighbours of an infected node. This parameter is relevant for this case study. Allen [6] identifies that the probability of a BBTV infection is inversely proportional to the distance from a previously infected plant, as most aphid flights cover small distances.
3. *Non-neighbouring probability of infection, θ*_*2*_: While short distance flights are more likely to occur in banana plantations, long distance aphid vector transmission remains a possibility. This parameter operates on all infected nodes, whereby each node has a probability of infecting every non-neighbour. Aphids are also known to be restless and sensitive to small changes in the environment and have been shown to relocate to other plants due to overpopulation, harvesting activities and sudden changes in atmospheric weather conditions [9].

The operations of these parameters are summarised in Fig 2 below:

**Fig 2.**
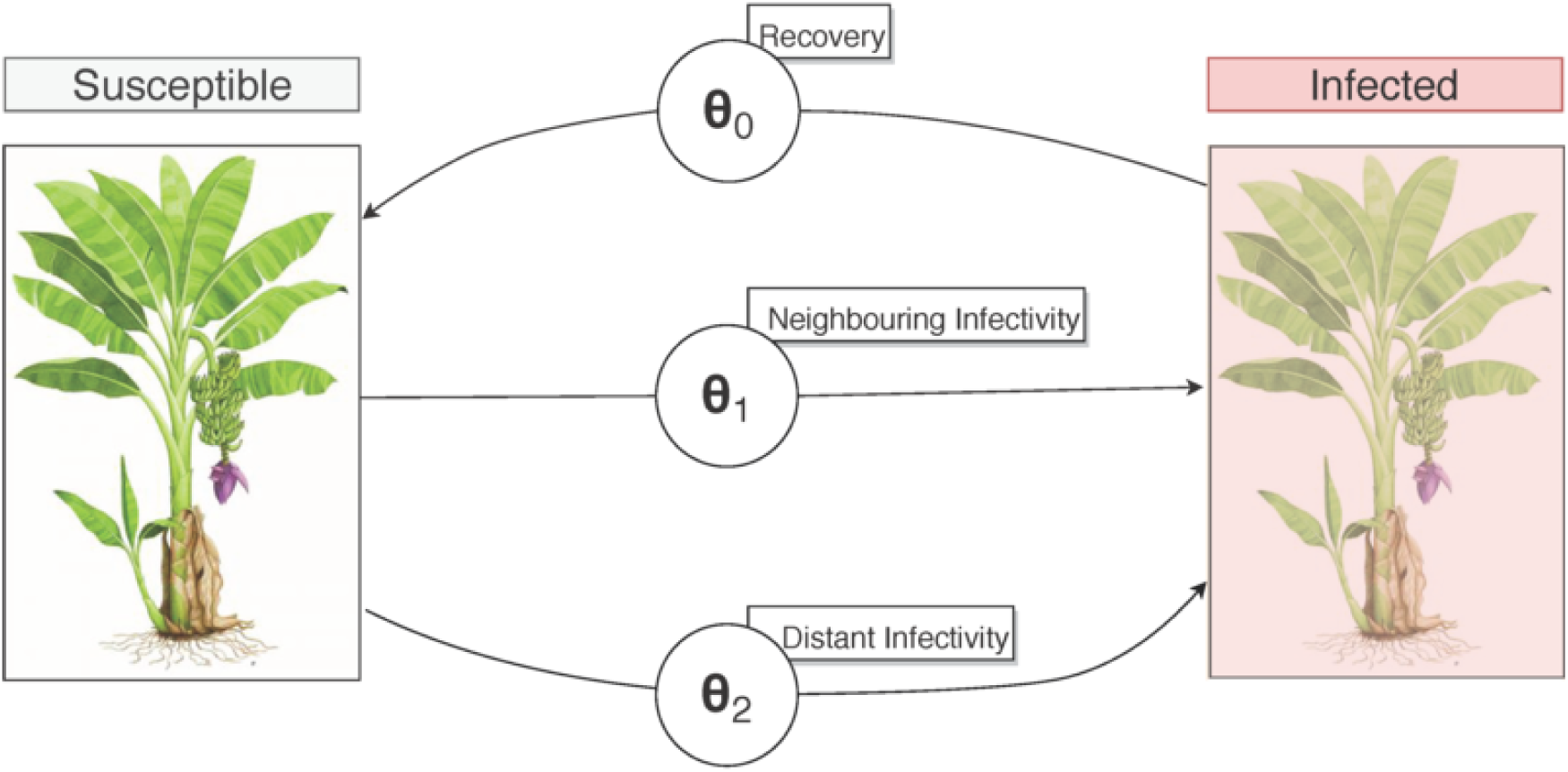
State-flow diagram describing each parameter’s influence on the state of a node. Banana plant has been chosen to represent an entire subsection of bananas.

#### Seasonality

BBTV spread is heavily influenced by seasonal changes in temperature, therefore it may be useful to identify changes in the posteriors for different seasons. Allen [6] identifies that detection efficiency, eradication efficiency and aphid activity are seasonally varying factors which greatly affect the spread of BBTV. Furthermore, Allen [10] confirms a seasonally varying leaf emergence rate in bananas, which informs detection and eradication efficiency. Anhalt and Almeida [11] observe temperature to be highly correlated with acquisition and inoculation efficiency, with peak transmission efficiency occurring at 25-30 degrees.

To accommodate for seasonal variation, we allow each parameter to be month-dependent. While it may be theoretically possible to generate unique posterior estimates for each month, their accuracy may be greatly diminished by the lower amount of field data available for each monthly parameter.

To maintain an effective sample size of observed data to inform each parameter while ensuring a clear differentiation between parameter counterparts, each parameter has been replicated by dividing and grouping months traditionally above or below the long-term average temperature. Months with an average temperature traditionally higher than the long-term annual average temperature, may be referred to as ‘summer’ months, and vice versa for ‘winter’ months. Summer months are given by September, October, November, December, January and February. Likewise, winter months are given by March, April, May, June, July and August.

Therefore, the final set of parameters are described by *θ*_*ij*_, where indicating the parameter type (recovery, near, and long distance infectivity respectively) and indicating the season (summer and winter respectively).

The mechanisms of the forward-simulation model are summarised in Fig 3.

**Fig 3.**
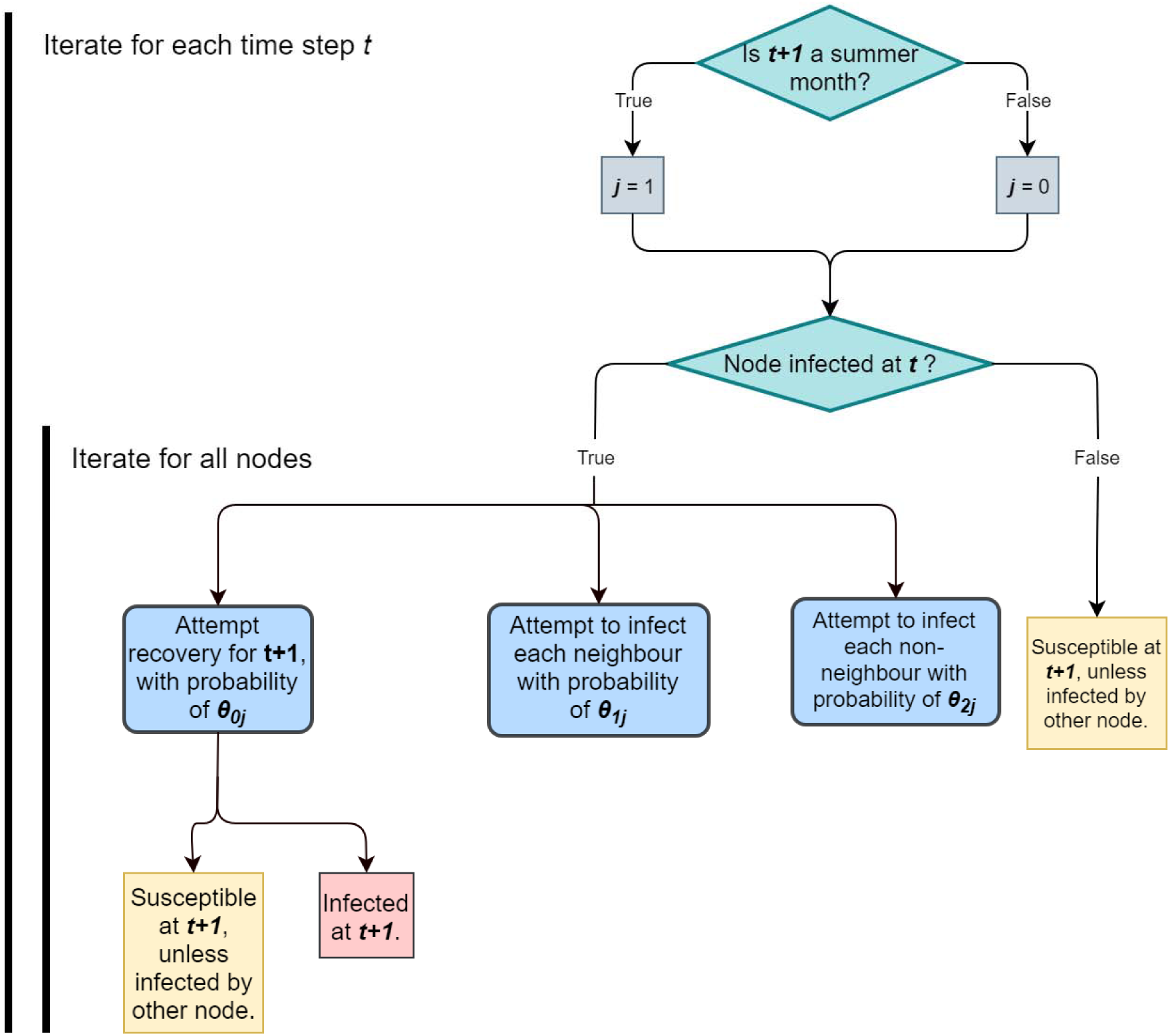
Flow-chart describing BBTV model behaviour for each time step *t*.

### Approximate Bayesian Computation (ABC)

The parameter *ϕ* = {*θ*_00_, *θ*_10_, *θ*_20_, *θ*_01_, *θ*_11_, *θ*_21_} may be inferred by its posterior density *p*(*ϕ* | *x*) given the observed dataset *x*= *x*_0_. The posterior density can be written by Bayes’ theorem as,

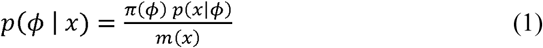

where *π*(*ϕ*), *p*(*ϕ* | *x*) and *m*(*x*) = ∫ *π*(*x*) *p*(*x*|*ϕ*) *dϕ* are, correspondingly, the prior density on the parameter *ϕ*, the likelihood function, and the marginal likelihood. The prior density *π*(*ϕ*) enables a way to leverage the learning of parameters from prior knowledge, such as the epidemiological characteristics and expert knowledge on BBTV [8]. As discussed in Dutta et al. [8], due to the complex stochastic nature of network disease simulator-based models (such as the one described in this paper), evaluating the likelihood function is computationally expensive while model forward simulation is relatively cheap. In this setting, approximate Bayesian computation [7] provides the opportunity to sample from the approximate posterior density of the parameters.

ABC bypasses the evaluation of the likelihood function by instead simulating data from the model to generate an approximate posterior distribution. Due to the high dimensionality of the observed data, *y*, the data set is often reduced to a set of summary statistics, *S*(*y*). Thus, ABC targets the posterior conditional on the summary statistics:

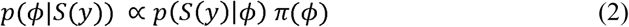

However, this too requires the evaluation of a typically intractable likelihood, *p*(*S*(*y*)|*ϕ*). Therefore, ABC approximates this intractable likelihood through the following integral:

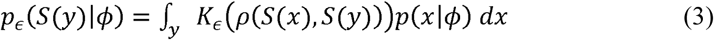

where *ρ*(*S*(*x*), *S*(*y*)) is a discrepancy function that compares the simulated and observed summary statistics, and *K*_*ϵ*_ (·) is a kernel weighting function with bandwidth *ϵ* that weights simulated summaries in accordance with their closeness to the observed summary statistic. The role of the discrepancy measure will become clear in the next section. While the integral in (3) is analytically intractable, it may be estimated by taking *n* iid simulations from the model 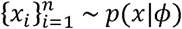, evaluating their corresponding summary statistics 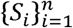 where *S*_*i*_ = *S*(*x*_*i*_), and calculating the following ABC likelihood:

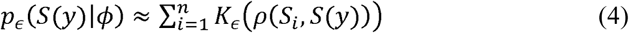

This unbiased likelihood estimator described in (4) is generally sufficient to obtain a Bayesian algorithm that targets the posterior distribution *p*_*ϵ*_(*ϕ*|*S*(*y*)) ∝ *p*_*ϵ*_ (*S*(*y*)|*ϕ*)*p*(*ϕ*). The summary statistics, *S*(·), discrepancy measure, *ρ*(·,·), and tolerance value, *ϵ*, utilised in the ABC method introduce approximation errors to the target posterior distribution. In order to minimise these errors, these factors must be chosen and tuned carefully to maximise accuracy while ensuring a computationally feasible operation [12].

#### ABC Algorithms

The most basic implementation of ABC is known as rejection sampling [13]. In this algorithm, the parameter *ϕ* is estimated by generating model realisations *x*_*sim*_ over the prior density space for *ϕ*. The summaries *S*(*x*_*sim*_) are computed and compared to *S*(*y*) through the discrepancy measure p(·,·). If the discrepancy between the simulated and observed summaries is lower than the tolerance, *ϵ*, then the proposed *ϕ*\* is accepted as part of the approximate posterior distribution.

The pseudo-code for an ABC rejection sampling scheme is provided below [14]:

~~~
*for i* ∈ 1: *n do*
        *Draw ϕ*\* ∼ *π(ϕ)*
       *Draw x*_*sim*_ ∼ *p*(·|*ϕ*\*)
      *Accept ϕ*\* *if p*(*x*_*sim*_, *y*) ≤ *ϵ*
*end for*
~~~

where *n* is the number of iid samples to be taken from the prior *π*(*ϕ*).

In this paper, ABC is implemented through a Markov Chain Monte Carlo (MCMC) algorithm to effectively estimate the posterior distributions of the recovery and spreading parameters of BBTV in the Newrybar banana plantation. ABC-MCMC [15] aims to improve the efficiency in comparison to ABC rejection sampling, by proposing parameter values locally around promising regions of the parameter space.

#### Summary Statistics

The summary statistics play an important role in the ability for ABC methods to effectively estimate the posterior distributions of parameters [12]. Summary statistics summarise observed or simulated data which can often be large, complex and high dimensional. Effective summary statistics characterise the influence of specific parameters on the model, so that varying parameter values result in observable changes in the reported summary statistics.

After some experimentation, we find that the following summary statistics are informative about the model parameters:

1. S_1_ – The proportion of infected nodes at each time step.
2. S_10_ – The total number of infected nodes at time *t*, that recovered by *t* + 1.
3. S_010_– The total number of susceptible nodes at time *t*, which became infected by *t* + 1. These nodes must not have an infected neighbour at time *t*.
4. S_011_ – The total number of susceptible nodes at time *t*, which became infected by *t* + 1. These nodes must have at least one infected neighbour at time *t*.

S_1_ describes the temporal characteristic of the simulated infection spread and is informed by all three parameters. S_10_ provides an indication of the recovery rate, and therefore informs θ_0_. S_010_ describes the number of infections occurring via a vector from a long distance, corresponding with θ_2_. S_010_ describes the number of infections occurring via a vector from a neighbouring subsection, informing θ_1_. Since the parameters are applied during different seasons, the summary statistics S_10_, S_010_ and S_011_ have been replicated for summer and winter. Given the time series data consists of 38 time points of which 20 are summer months, we have a total of 44 summary statistics.

#### Discrepancy Measure

The Mean Squared Error (MSE) between the simulated and observed summary statistics was utilised as a discrepancy measure for this case study.

#### Priors

Given the novel nature and the lack of specific expert knowledge regarding these parameters, uniform priors with bounds of [0,1] have been chosen for all parameters.

## Results and Discussion

### Posterior Distributions

Mean posterior recovery probability (*θ*_*0j*_) is significantly influenced by the seasonal changes in temperature. While the mean posterior recovery probability for an infected subsection is 25.7% in summer (θ_01_), this increases to 30.63% for winter (*θ*_*00*_) (Fig 4).

**Fig 4.**
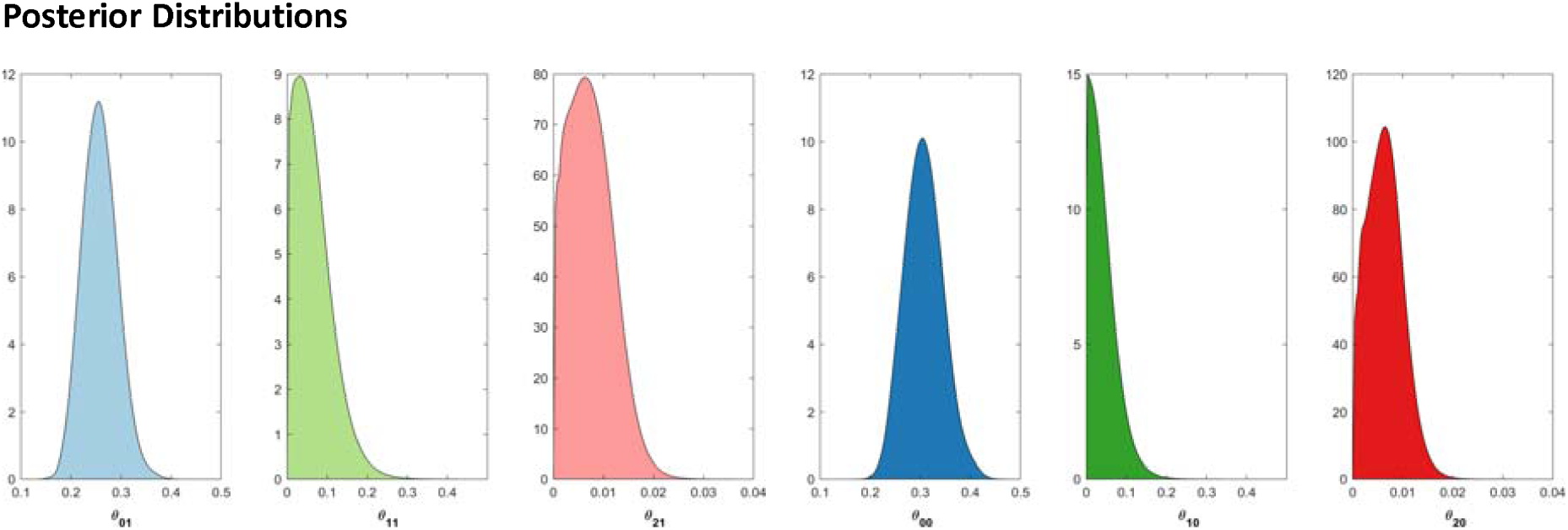
Posterior distributions of parameters (1e7 MCMC iterations of which 1e5 MCMC burn-in) Posterior densities are coloured to correspond with their respective seasonal counterpart, with darker colours representing winter seasons.

Mean posterior neighbouring infectivity (*θ*_*1j*_) is 2% higher in summer months compared to winter months (6.2% and 4.1% respectively), indicating minor seasonal dependence. Distant infectivity exhibits a similar dependence (*θ*_*2j*_), as the mean posterior probability during summer of 0.07% only decreases by 0.01% during winter months (Fig 4).

These results may be explained through a range of environmental factors. Higher posterior probabilities for node recovery in winter are likely due to lower inspection accuracy arising from lower leaf growth rates and decreased farming activity during this season. Research from the Department of Employment Economic Development and Innovation identify that bananas growing in the sub-tropical climates such as the South-East Queensland area are heavily influenced by temperatures: the rate of production is often significantly reduced in winter, sometimes to a rate of one leaf in 20 days [16]. In contrast, summer leaf emergence can be completed in around four days in tropical conditions. Since BBTV in banana plants is identified through observing visual symptoms of infections, identifying newly infected plants is much more likely during summer compared to winter, where a plant may be latently infected for months before displaying signs of infection. Therefore, the lower reported BBTV infection rates in winter would artificially increase the posterior probability of recovery in this season.

Higher neighbouring and distant infectivity in summer is likely due to more weather events in this season, and inoculum acquisition sensitivity to tropical temperatures. According to the historical monthly averages of climate data provided by the Bureau of Meteorology, South-east Queensland experiences significantly higher wind speeds during summer at an average maximum gust of 131.7 km/h compared to 93.7 km/h during winter [17]. Similarly, average rainfall during summer is 113.7 mm for 11 days compared to 84.2 mm for nine days during winter [17]. A higher frequency and intensity of such weather events are likely to perturb the aphid vector, as observed by Claflin et al. [9], increasing the risk of neighbouring and long-distance transmission. Additionally, Anhalt & Almeida [11] identify that *P. nigronervosa* provides peak inoculum acquisition and transmission efficiency between 25-30 degrees Celsius, which is historically experienced during the summer months in South-east Queensland. Higher seasonal rates of inoculum transmission, in conjunction with increased vector activity during this period could account for greater counts of neighbouring and long-distance infections during the summer.

The estimated pairwise posterior distributions of model parameters highlight key operative characteristics of the model. As seen in Fig 5, the estimated univariate posterior distribution for the posterior probability of recovery is largely symmetric, while the posterior probability of neighbouring and distant infectivity (*θ*_*1j*_) remain positively skewed.

**Fig 5.**
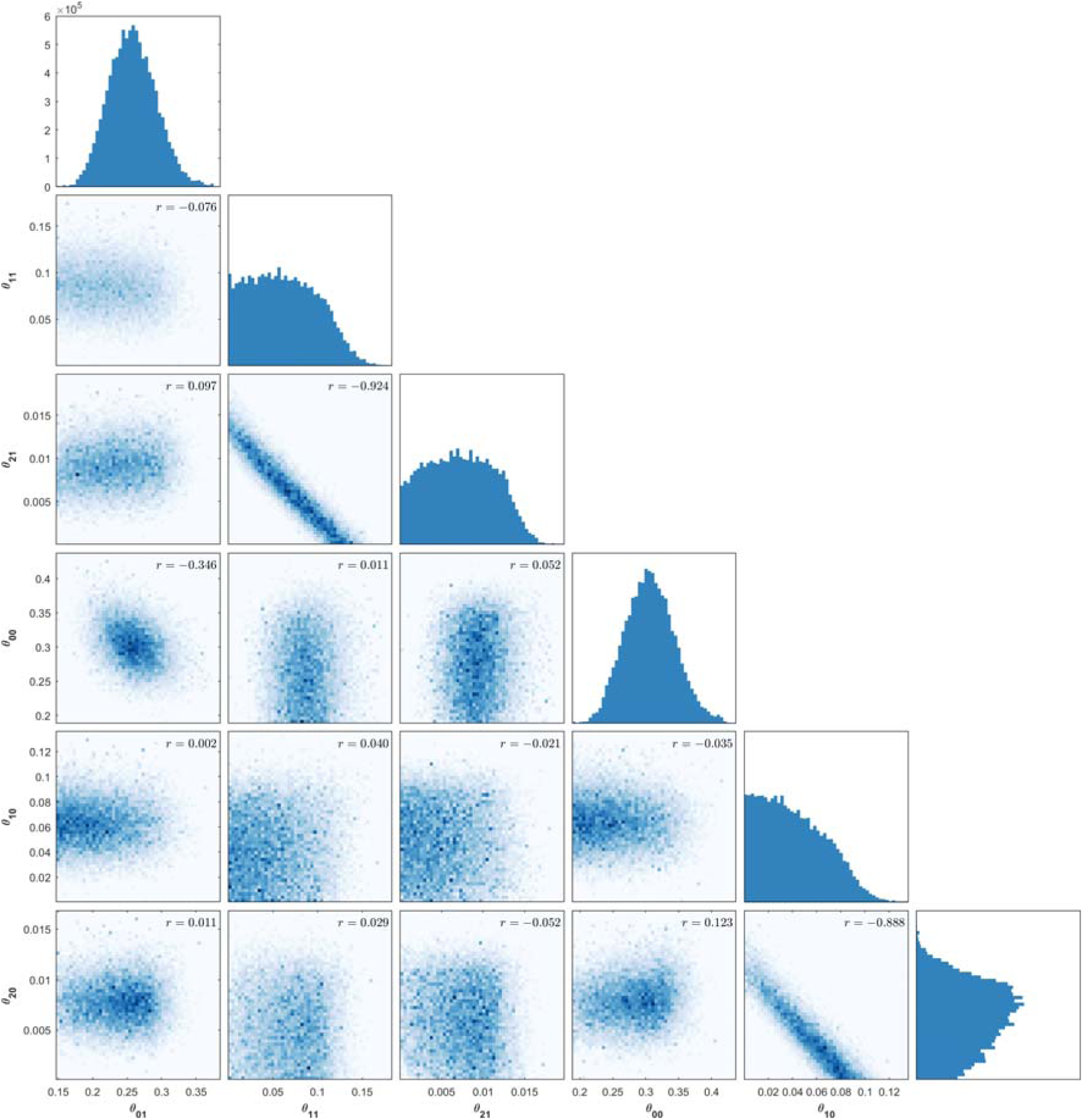
Estimated univariate and pairwise posterior distributions using MCMC-ABC.

Negative correlation is evident between neighbouring infectivity (*θ*_*1j*_) and distant infectivity in the corresponding months (*θ*_*2j*_), which indicates that different combinations of these parameters can generate similar summary statistics. This is an intuitive correlation since the total number of infected nodes in each month is a sum of the number of nodes infected by a neighbour or over long-distance. Therefore, if a high probability of neighbouring infectivity is proposed in a model simulation, a low probability of distant infectivity will be more likely to result in the observed summary statistics.

### Posterior Forecasting

Fig 6 provides a posterior infectivity forecast for all subsections over a 6-month period. The forecast is generated by running the BBTV forward simulation model with parameters obtained from a random sample from the posterior distribution.

**Fig 6.**
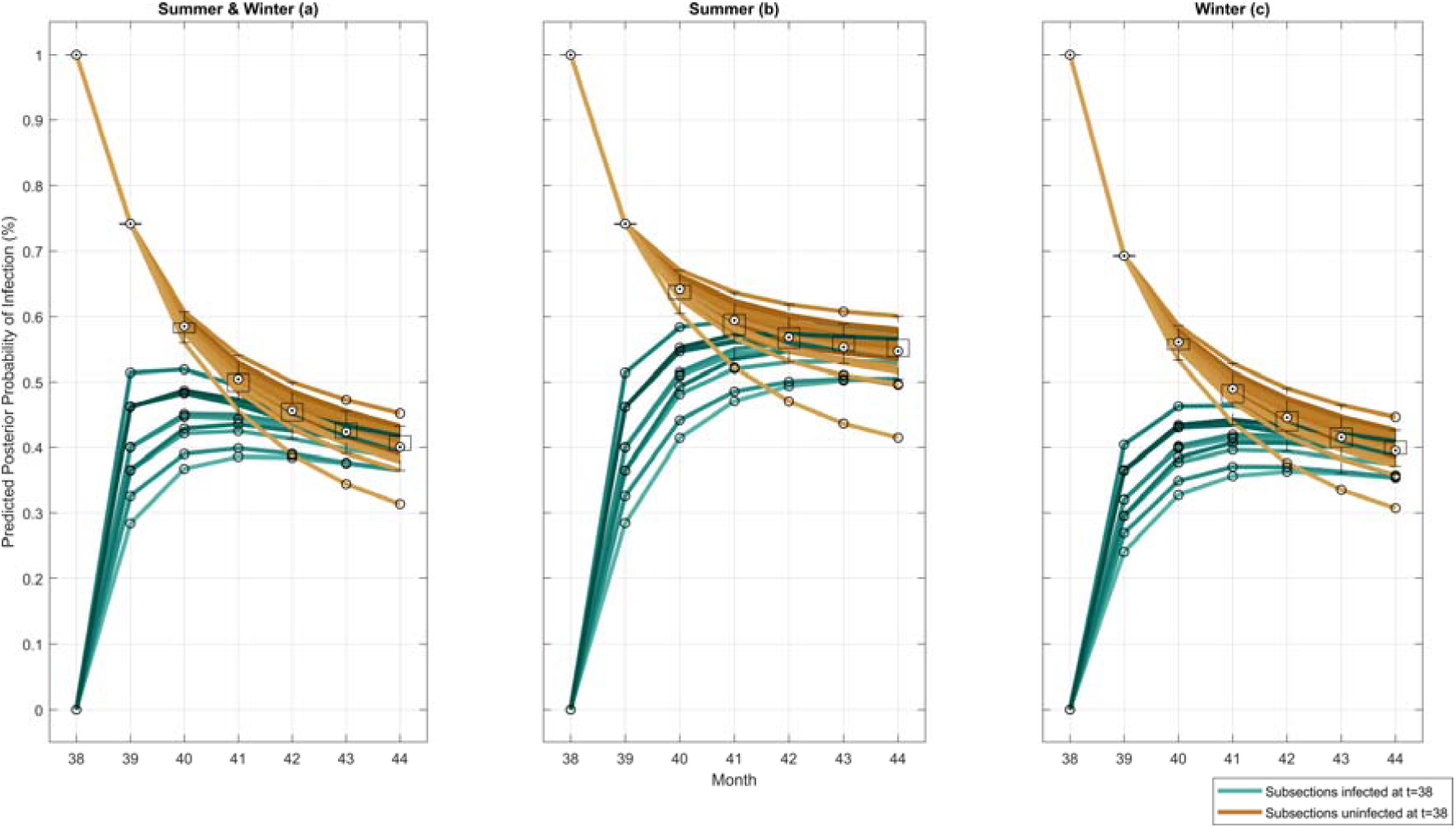
6-month posterior forecast of subsection infectivity. (a) Simulated using both summer and winter posterior counterparts. (b) Simulated using only summer posterior counterparts. (c) Simulated using only winter posterior counterparts.

Each node begins with a 100% or 0% infection probability in month 38, since this is the last known time-step provided to the model as the initial configuration for each of the three simulation scenarios. The subsequent forecasted probabilities of infection, for each plantation subsection, converge to a steady state probability of 45%. The forecasted infection probabilities for subsections infected in the last known time-step are characterised by a depreciation in infection probability, and an increasing variance in individual infection probability at each subsequent month. In contrast, forecasted probabilities for initially uninfected subsections begin with a high variance in the group, progressively decreasing in subsequent months.

Fig 6(b) provides a posterior forecast of subsection infectivity which applies the posterior counterparts for summer months to all months (*θ*_*i1*_). Compared to Fig 6(a), this results in a higher steady state probability of 57%. Furthermore, while previously uninfected subsections in both figures exhibit a steep increase in forecasted subsection infectivity by the first month, the subsequent infection probabilities flatten to the steady state infection probability in Fig 6(a) and continue to increase in Fig 6(b). This is likely due to the higher neighbour infectivity and distant infectivity probabilities during summer.

This may be confirmed through Fig 6(c), which provides a 6-month posterior forecast of subsection infectivity, utilising the posterior counterparts of winter months to all months (*θ*_*i0*_). Unlike Fig 6(b), Fig 6(c) displays a gradual increase in forecasted probability of subsection infectivity over subsequent months, tending to a lower steady-state subsection infectivity probability of 40%.

### Discrete-space posterior probability forecast

Fig 7(a) and 7(b) visualise the 1-month posterior probability forecasts for infection in each plantation subsection. Fig 7(a) depicts the posterior infection probabilities for infected subsections in the last observed time-step, while 7(b) depicts the posterior infection probabilities for previously uninfected subsections.

**Fig 7.**
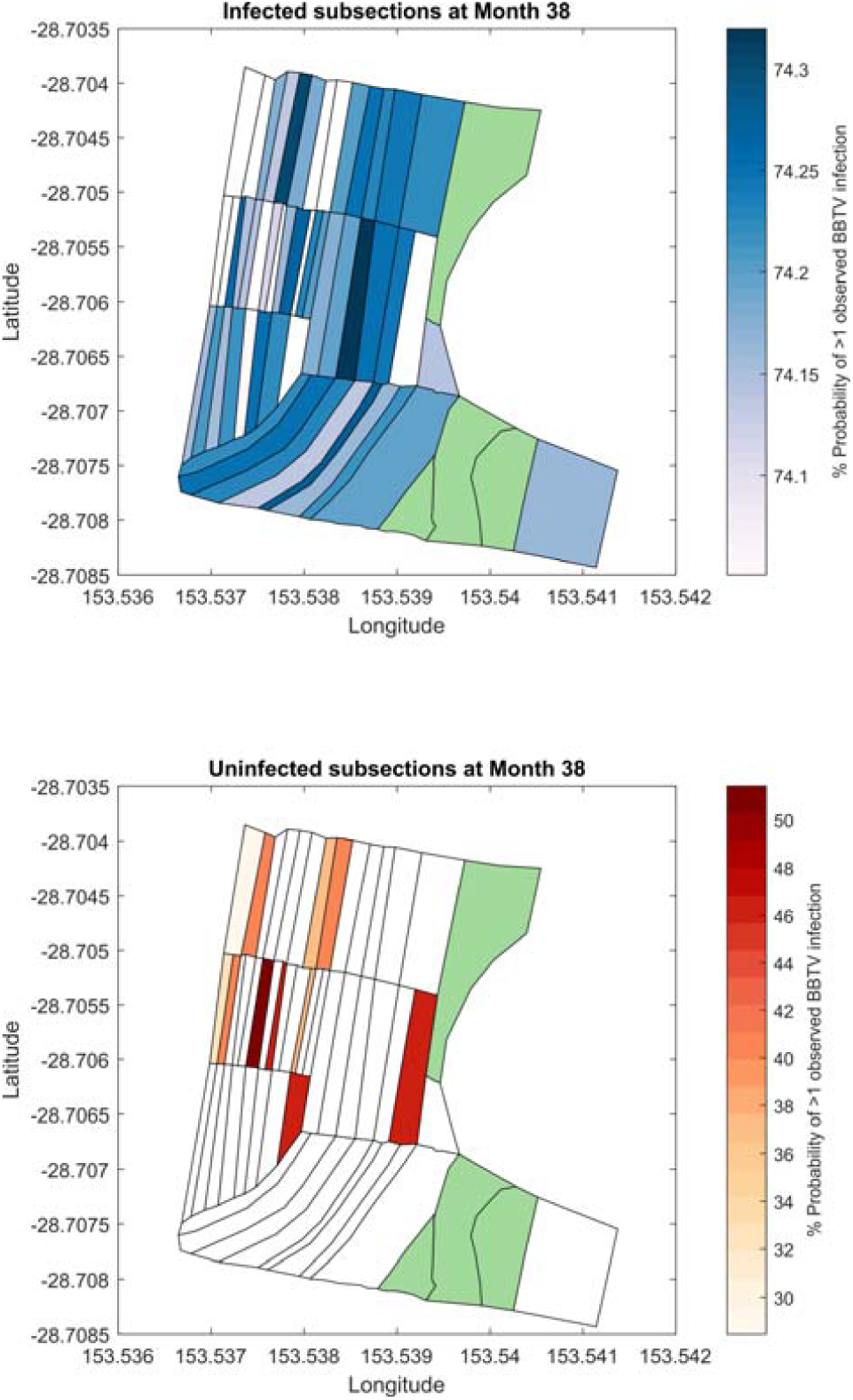
1-month discrete-space posterior forecast of subsection infectivity. (a) Posterior probability of subsection infection for previously infected subsections. (b) Posterior probability of subsection infection for previously uninfected subsections. Subsections highlighted in green indicate areas not planted with bananas.

Posterior forecasts from the BBTV model indicate that previously infected subsections are likely to remain infected, consistently reporting an infection probability of approximately 74% for all subsections. Since the forecasted month (39) occurs during the summer period, the posterior distributions of the summer counterparts (*θ*_*i1*_) are utilised for this simulation.

Figs 7(a) and 7(b) indicate that the highest posterior predicted infection probability is associated to subsections with a high number of neighbours. This is likely due to the greater application of neighbouring infectivity (*θ*_*1j*_) on this subsection, as a high number of neighbours increases the probability of the aphid vector to infect this subsection.

### Alternate Plantation Organisation

The model may be utilised to explore the reconfiguration of a banana plantation, while removing areas of the plantation might reduce the infection probability of the subsection. Fig 8(a) and 8(b) describe the 1-month posterior infection probability forecasts for month 39 if subsections 20 to 34 (in grey) were cleared.

**Fig 8.**
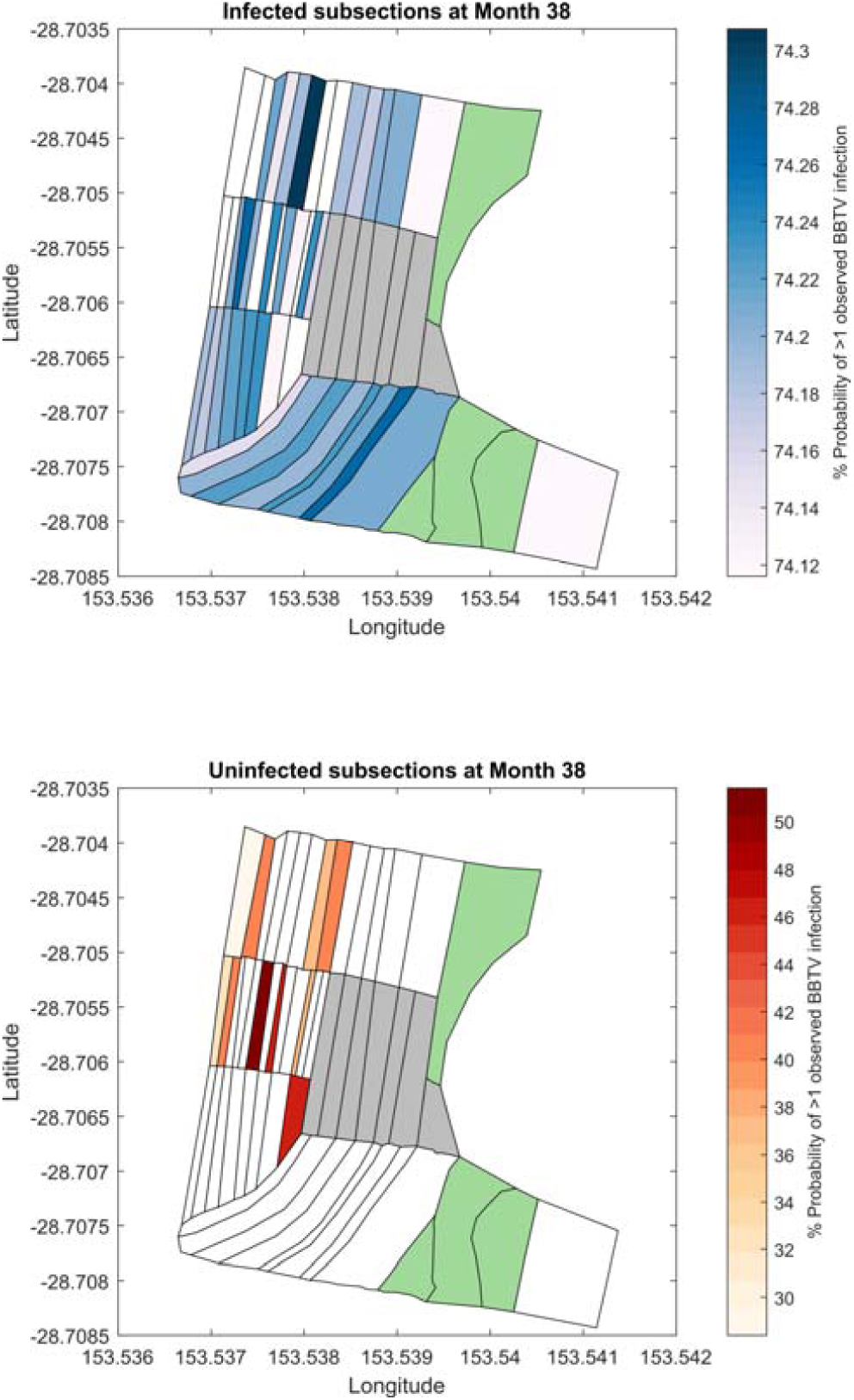
1-month discrete-space posterior forecast of subsection infectivity. (a) Posterior probability of subsection infection for previously infected subsections. (b) Posterior probability of subsection infection for previously uninfected subsections. Subsections highlighted green indicate areas not planted with bananas. Subsections highlighted grey indicate areas that have been ‘cleared’ according to the simulation.

When compared to the forecasts from Fig 7(a) and 7(b), removing subsections 24 to 32 results in an insignificant reduction in forecasted posterior infection probabilities by approximately 0.4% for each subsection. Further configurations of cleared areas may be explored upon recommendations by stakeholders.

### Implications and Future Research

Monitoring the nodal recovery (*θ*_0*j*_), near infectivity (*θ*_1*j*_) and distant infectivity (*θ*_2*j*_) rates over time would serve to metricise the effectiveness of disease management strategies. For example, a long-term reduction in *θ*_*0j*_ could be a signal to increased aphid resistance to chemicals used in current pesticide routines, or an increase in latent infections across the plantation. Correspondingly, a long-term increase in *θ*_*1j*_ and *θ*_*2j*_ could point to higher aphid activity across the plantation, potentially indicating a reduction in inspection accuracy.

Providing site inspectors with specific high-risk areas through posterior predictive modelling could improve their inspection accuracy, as it would support their priori of likely sites of infection across the plantation. Furthermore, the identification of 3 or 4 high-risk areas in a plantation opens the opportunity of increasing the site inspection frequency from monthly inspections to fortnightly inspections, while limiting the inspection coverage to these high-risk areas.

The model has several limitations. Firstly, the model does not account for the geographical characteristics of the plantation. The steepness and height of an area of the plantation would affect the ability for the aphid vector to travel to neighbouring subsections. Furthermore, the area of each subsection, and the distance to neighbouring subsections is quite varied and would be likely to influence the posterior probability of neighbouring and distant infectivity due to the greater distance for the aphid vector to travel, and the number of potential host plants in a subsection. Secondly, environmental factors such as the wind speed and direction, and extreme weather events are not considered in this model. Higher wind speeds and extreme weather events would be more likely to perturb and relocate the aphid vector, resulting in greater infectivity rates across the plantation. Thirdly, the methodology places limits on the achievable resolution of posteriors, since the model relies on the presence of clear divided areas in a banana plantation to identify subsections and establish a network. Furthermore, since the collection of field data relies on visual observation, the accuracy of model parameters largely relies on the inspection accuracy which is influenced by seasonality and environmental factors, in addition to inspector experience and fatigue. Lastly, the model only considers the presence of at least one infection in a subsection, whereas the field data collected at Newrybar would enable the calculation of infection counts in each subsection, as well as the total number of infected leaves present in a subsection. These factors would significantly improve model accuracy and informativeness. Extending the model to address these limitations should be considered for future research.

## Conclusion

This paper has adapted and extended upon current network-based disease models implemented in an ABC framework. A forward-simulating network-based SI model has been created which simulates the spread of BBTV across the subsections of a banana plantation, by parametrising nodal recovery, neighbouring infectivity and distant infectivity across summer and winter. Findings from posterior results achieved through MCMC-ABC indicate significant seasonality in all parameters, which are influenced by correlated changes in inspection accuracy, temperatures and aphid activity. This model enables the simulation, monitoring and forecasting of various disease management strategies, which may support policy-level decision making and inspector experience. Introducing higher dimensional field and weather data will improve model accuracy and utility; an area to be explored for future research.

## Acknowledgements

The authors would like to acknowledge the support of Hort Innovation in providing the data required for this project, and Barry Sullivan, Dr. John Thomas and Dr. Kathy Crew for their expert advice on BBTV throughout this project.

## Supporting information

### Posterior Predictive Checking

In addition to the 38 months of field data provided by the BBTV Prevention Program from December 2014 to January 2018, an extra 7 months of field data till August 2018 was provided, which may be utilised as validation dataset. In order to conduct posterior predictive checking, the model was set to simulate the posterior infection probabilities of each node in month (*t* + 1), given an initial configuration of the infected nodes at month *t*. The validation data for month (*t* + 1) may then be compared to the corresponding posterior infection probabilities to identify model accuracy.

#### Binomial Deviance Loss

The accuracy of the posterior predicted infection probabilities for a subsection may be identified through its deviance from the validation data. This may be calculated as follows:

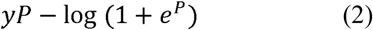

where *y* is the actual data for a subsection in month (*t* + 1), and *P* is the log odds of the corresponding posterior prediction.

Fig S1 displays the binomial deviance loss for each prediction for each subsection, coloured according to the validation dataset used. Prediction accuracy is largely unaffected by the validation set used, with the mean deviance loss across all subsections being -0.6; a 40% improvement compared to a random prediction. Posterior predictions for certain subsections are consistently excellent (see subsections 27-32, 43-50), due to consistent incidence levels in these areas of the farm.

#### Receiver-Operating Characteristic (ROC) Curve

A ROC Curve illustrates the diagnostic ability of a binary classifier system as its discrimination threshold is varied. The ROC curve is created by plotting the true positive rate (TPR) against the false positive rate (FPR) at various threshold settings. Fig S2 displays the ROC curve of the model, which demonstrates an Area-Under-the-Curve (AUC) of 0.65, generally considered to be a “fair” model. This metric is a general indicator of the posterior prediction confidence, as a larger area under the curve would indicate higher true positive rates at lower threshold levels. The ROC curve is well above the diagonal line, indicating that it performs markedly better than random predictions, particularly at higher thresholds.

---

**Fig S1:**
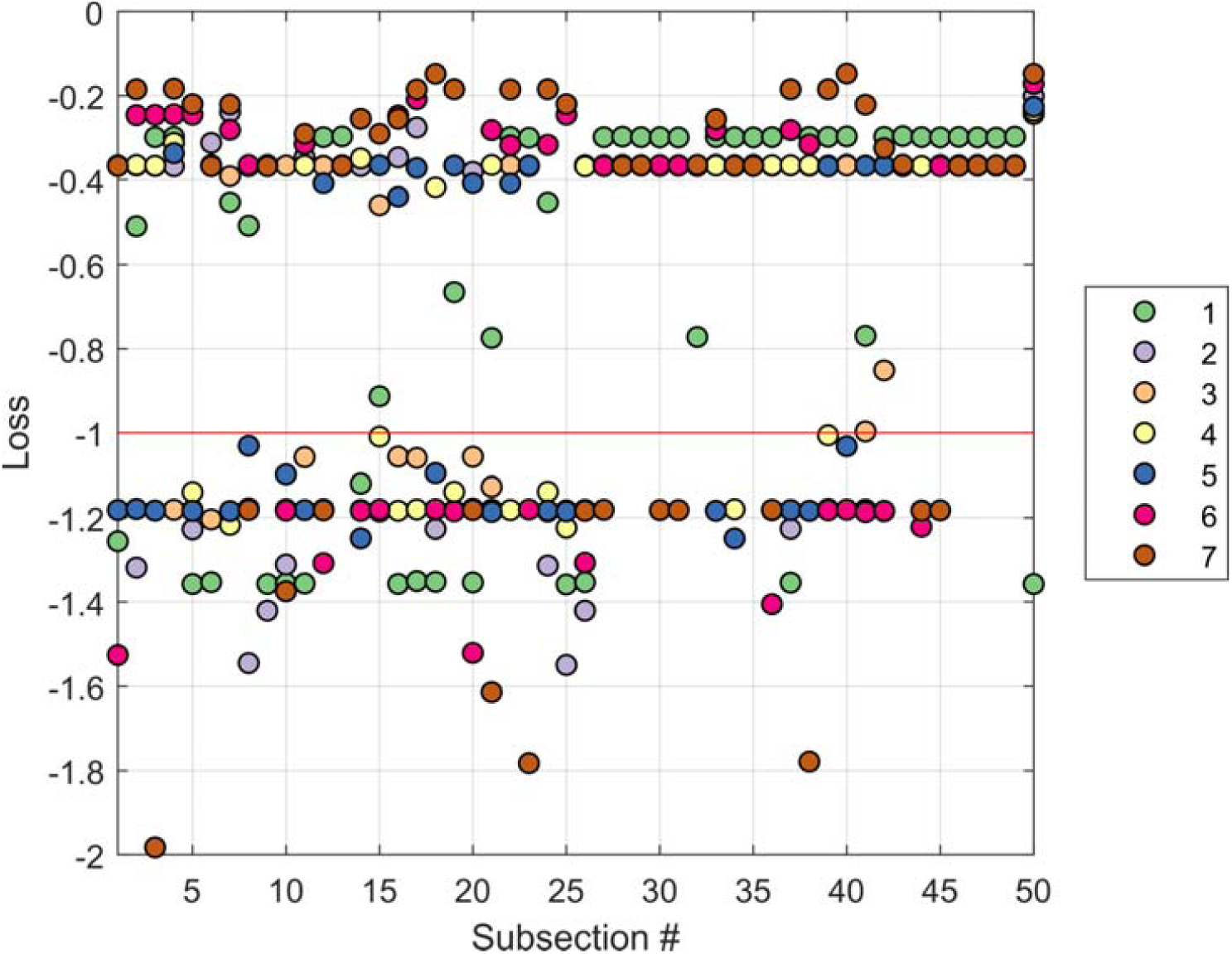
Binomial Deviance Loss aggregated for all predicted months in a 6-month period. Each dot represents the binomial deviance for a prediction corresponding to a subsection, coloured according to the predicted validation data set. The red line represents the deviance loss of a posterior prediction of 50% (random).

**Fig S2:**
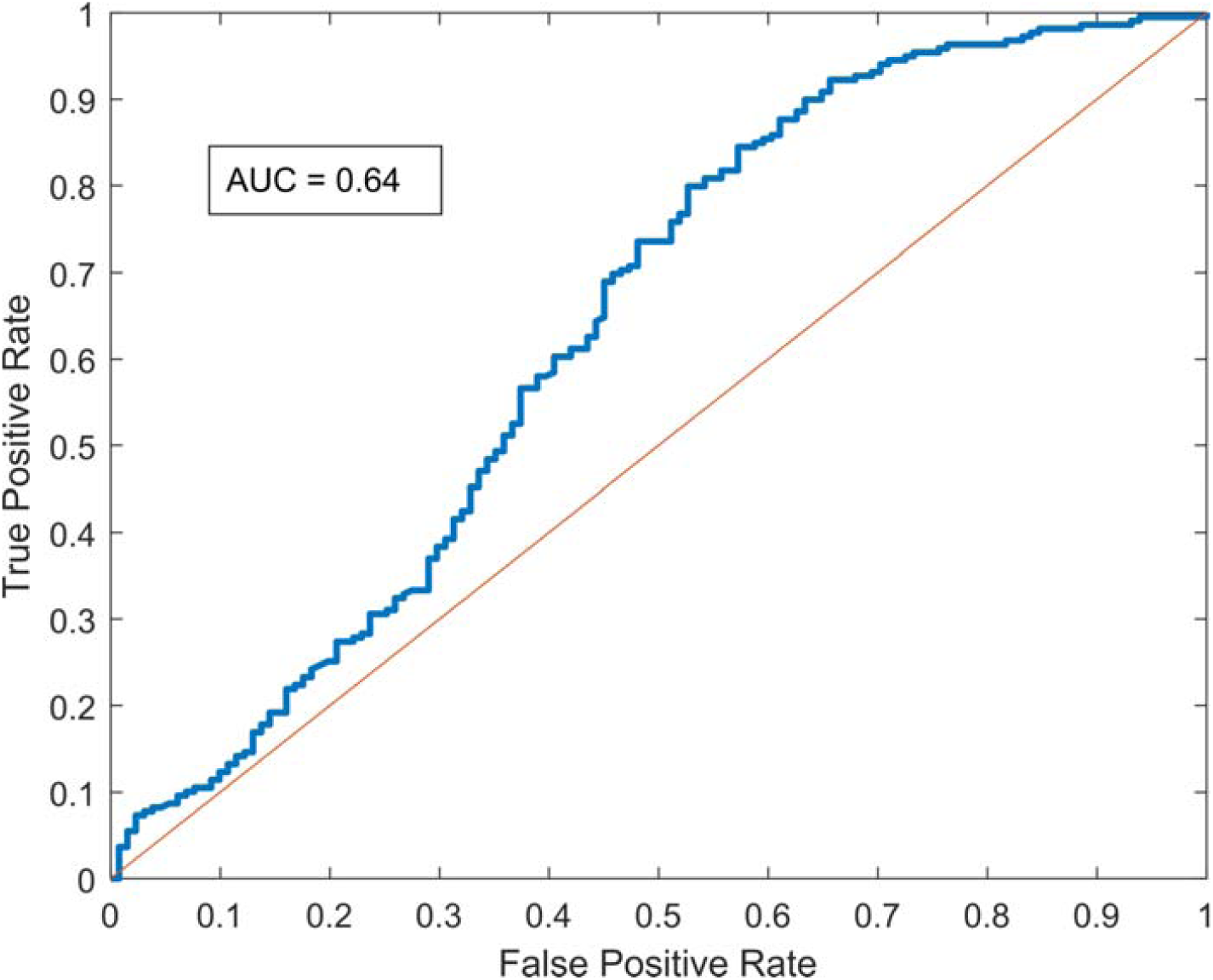
Receiver-Operating Characteristic (ROC) Curve for the BBTV forward-simulating model. Predictions across all validation data sets have been aggregated to construct this curve.

## References

1. Smith MC, Holt J, Kenyon L, Foot C. Quantitative epidemiology of Banana Bunchy Top Virus Disease and its control. Plant Pathology. 1998;47(2):177–87.

2. Fish S. The History of Plant Pathology in Australia. Annual Review of Phytopathology. 1970;8(1):13–36.

3. Cook DC, Liu S, Edwards J, Villalta ON, Aurambout J-P, Kriticos DJ, et al. Predicting the Benefits of Banana Bunchy Top Virus Exclusion from Commercial Plantations in Australia (Banana Bunchy Top Virus Control in Australia). 2012;7(8):e42391.

4. Dale JL. Banana bunchy top: an economically important tropical plant virus disease. Advances in virus research. 1987;33:301.

5. Brooks-Pollock E, Roberts GO, Keeling MJ. A dynamic model of bovine tuberculosis spread and control in Great Britain. Nature. 2014;511:228.

6. Allen RN, New South Wales Department of Agriculture WNCAI. Further studies on epidemiological factors influencing control of banana bunch top disease, and evaluation of control measures by computer simulation. Australian Journal of Agricultural Research; ISSN. 1987;38(2):373–82.

7. Sisson SA. Handbook of Approximate Bayesian Computation. Fan Y, Beaumont M, editors. Milton: CRC Press LLC; 2018.

8. Dutta R, Mira A, Onnela JP. Bayesian inference of spreading processes on networks. Proceedings Of The Royal Society A-Mathematical Physical And Engineering Sc. 2018;474(2215).

9. Claflin SB, Power AG, Thaler JS. Aphid density and community composition differentially affect apterous aphid movement and plant virus transmission. Ecological Entomology. 2017;42(3):245–54.

10. Allen RN, New South Wales Department of Agriculture WARC. Epidemiological factors influencing the success of roguing for the control of Bunchy Top disease of bananas in New South Wales. Australian Journal of Agricultural Research; ISSN. 1978;29(3):535–44.

11. Anhalt MD, Almeida RPP. Effect of Temperature, Vector Life Stage, and Plant Access Period on Transmission of Banana bunchy top virus to Banana. Phytopathology. 2008;98(6):743–8.

12. Dennis P. Summary Statistics. Handbook of Approximate Bayesian Computation: CRC Press; 2018.

13. Beaumont MA, Zhang W, Balding DJ. Approximate Bayesian Computation in Population Genetics. Genetics. 2002;162(4):2025.

14. Marin J-M, Pudlo P, Robert CP, Ryder RJ. Approximate Bayesian computational methods. Statistics and Computing. 2012;22(6):1167–80.

15. Marjoram P, Molitor J, Plagnol V, Tavaré S. Markov chain Monte Carlo without likelihoods. 2003;100(26):15324–8.

16. Deuter P, White N, Putland D, Mackenzie R, Muller J. Critical (temperature) thresholds and climate change impacts/adaptation in horticulture; 2011.

17. Ballina Weather Station: Bureau of Meteorology. In: Meteorology Bo, editor. [online]: Ballina Airport Weather Station; 2018.

